# Anti-HIV-1 Activity of *Crocodylus mindorensis* (Philippine crocodile) serum in cell-free and cell-associated virus interactions to human peripheral blood mononuclear cells

**DOI:** 10.1101/596775

**Authors:** Alfredo Hinay, Nelyn Mae Cadotdot, Marilou Tablizo

**Author notes:** Corresponding author: Alfredo Jr, Hinay, Faculty of Medical Laboratory Science, University of the Immaculate Conception, Davao City, Philippines.

## Abstract

Highly-Active Antiretroviral Therapy (HAART) is the recommended treatment and management strategy for HIV infection. Although the existing antiretroviral drugs are indispensably significant in improving the quality of lives of HIV/AIDS individuals, the drugs still have many limitations including resistance, production of toxicity, and their limited availability. These limitations continue to open new opportunities in the use of ethnomedicine for the management of HIV/AIDS. With this, few researchers have made an effort to test the inhibitory activity of crocodile serum as it has a unique and diverse molecular activity in preventing HIV-1 replication. In this study, a cell culture-based assay was utilized coupled with colorimetric enzyme immunoassay to determine the HIV-1 reverse transcriptase activity. One HIV-1 seropositive serum was processed for Peripheral Blood Mononuclear Cells (PBMC) co-culture from which HIV-1 isolates were obtained. The HIV-1 reverse transcriptase activity after 21 days was 0.5928 pg/well. Moreover, a baseline Philippine crocodile serum concentration of 0.5% vol/vol was used based on the previous study conducted by Hinay and Sarol (2018) and the cell viability results showed no cell reduction of mononuclear cells after 72 hours incubation. The inhibitory activity of the Philippine crocodile serum at 0.5% and 0.25% vol/vol concentrations inhibited 65.68±2.93% and 69.92±0.45% respectively in post-infection interactions. In addition, the Philippine crocodile serum in pre-infection interaction at 0.5% and 0.25% vol/vol concentrations inhibited 68.61±1.67% and 69.95±2.24% respectively. As has been noted, the inhibitory actions of the Philippine crocodile serum effectively regulate the HIV-1 replication in both pre- and post-infection interactions.

## Introduction

Reptiles such as alligators exhibit remarkable ability to heal rapidly considering the septic environment in which they live. Merchant and colleagues provided observation on the broad range of antiviral activity of alligator serum (Merchant et. al, 2003, 2004). Merchant’s recent studies also showed that these activities are due to a potent and broad-acting serum complement system (Merchant et. al, 2004). In the Philippines, there are two species of crocodile – the *Crocodylus porosus* locally known as Philippine saltwater crocodile and the *Crocodylus mindorensis*, the endemic Mindoro crocodile. The relatedness of *Alligator mississippiensis* (*American alligator*) described by Merchant and colleagues and the Philippine crocodile *Crocodylus mindorensis* based on phylogeny shows that they belong to the same phylum (Crocodilia) but from different Families (Casey, Gardner, & Farke, 2012).

Crocodiles may acquire serious injuries when they hunt food and may encounter higher order animals to compete for survival. However, the crocodile’s immune system responds well to these injuries and the crocodiles generally survive and do not show signs of illness. Moreover, crocodilians have been shown to have a strong immune system with a remarkably higher biological and pharmacological activity than others animals, and also humans, which make them a good platform for drug discovery (Dzik, 2010). These immunological effector mechanisms necessary for the efficient control of infectious agents such Human Immunodeficiency Virus Type-1 that are dependent on distinct defense strategies. Some components involved in these routes of the defense system have been identified and characterized which probably include serum complement cascades (Merchant et al., 2003; Dzik, 2010), serum Mannose-Binding Protein (Ezekowitz, Kuhlman, Groopman, & Byrn, 1989) and conglutinin-like protein (Ushiijima, et al., 1992). All these features may contribute to the antimicrobial and antiviral properties of crocodile serum.

The antiviral activity of *Alligator mississippiensis* (American alligator) serum described by Merchant, 2005 conducted using cell culture-based method provided the evidence that it contains anti-HIV-1 activity with half maximal inhibitory activity of 0.9% (Merchant et. al, 2003, 2004). In the Philippines, particularly in Davao, crocodiles which are characteristically similar to alligators are not only abundant but are scientifically cultured and propagated. With the availability of Philippine crocodile, a scientific research that will be a foothold and basis that *Crocodylus mindorensis* serum could be used as a potential inhibitor of HIV/AIDS infection was investigated.

## Subjects and Methods

### Specimen Collection

A minimum of 2 mL serum sample from one (1) purposively selected adult-male *Crocodylus mindorensis* was collected at the Animal Clinic of the Davao Crocodile Park facility. The freshly collected sera were separated from whole blood samples collected by a veterinarian as part of the health check of the crocodile. Specifically, 5 mL of whole blood were allowed to clot at room temperature for approximately three (3) hours. The serum was separated by centrifugation at 3,000 × g for 15 minutes and processed immediately.

### Ethical Considerations

#### Blood Sampling Protocol

Peripheral blood mononuclear cells were obtained from two different individual sources. First from a healthy non-HIV-1 reactive individual and second from a HIV-1 reactive individual. An inform consent were given to both participants before blood collection.

A volunteer HIV-1 reactive individual with a signed inform consent was the source for HIV-1 co-culture method. A three (3) 5 mL EDTA tube was used to collect the blood using venepuncture technique with strict compliance to Occupational Safety and Health Administration 1910.1030 Bloodborne Pathogens guidelines (OSHA 1910.1030). After collection, the sample was processed immediately. The same protocol was followed for healthy non-HIV-1 reactive volunteer.

#### Biosafety Considerations

Upon arrival of crocodile serum sample, proper protocol was observed including serum separation to anti-HIV-1 activity assessment. At the end of the experiment, the used materials such as microplate and washing basin containing 5.25% sodium hypochlorite were decontaminated at 121°C at 15psi for 30 minutes including all crocodile serum received.

#### HIV-1 Isolation

Isolation of HIV-1 from one sero-positive sample was done using a PBMC micro-co-culture assay (Dahake *et.al* 2013). Briefly, one healthy, HIV-sero-negative donor PBMCs were stimulated with the mitogen phytohemagglutinin-P (PHA-P; at a final concentration of 5.0 μg/mL), in the presence of human interleukin 2 for 24-72 hours before use to promote blast formation and replication of T-cells. The cells were counted and 1×10^6^ PHA-stimulated donor cells and 1×10^6^ PBMCs from sero-positive HIV-1 individual was added in duplicate wells of a 24-well tissue culture plate. The final volume was adjusted to 2.0 mL with growth media containing RPMI1640 with 20% FBS and 10.0 U/mL IL-2. The plate was incubated at 37°C with 5% CO2 in a CO2 incubator. On day 7 and 14, the cultures were replenished with fresh growth media containing 5×10^5^ PHA stimulated donor cells (feeder cells). The culture was continued until day 21 and then terminated. Supernatant fractions from duplicate wells from day 14 and 21 were saved separately, and stored at −70°C until analysis for HIV-1 RT by ELISA. Cultures were considered positive only when day 21 supernatants showed an increase in HIV-1 RT from day 14 supernatants.

### Cell viability assay

Before using the Philippine crocodile serum sample for antiviral assay, it was necessary to assess their toxicity against Peripheral Blood Mononuclear Cells (PBMC). Briefly, a 100 uL PBMC was suspended into a 0.5 mL microcentrifuge tube and added 90 uL of Safranin stain. The PBMC were counted to 1×10^6^ cells. After determining the PBMC count, cell viability assay was performed. Briefly, a 100 uL 1×10^6^ PBMC were suspended into a 0.5 mL microcentrifuge tube and added 100 uL of 0.5% and 0.25% vol/vol Philippine crocodile serum. The interactions were allowed up to 72 hours and cell viability was counted.

#### Post-infection interaction (cell-associated HIV)

The procedure was based on the study of Dahake, R (2013). Briefly, HIV-1 virions from co-cultured were first allowed to infect PBMCs and the Philippine crocodile serum were added to the suspension after one hour. A volume of 100 μL of HIV-1 virions were added to 1 mL of PHA-stimulated PBMCs and incubated for 2 hours in a CO_2_ incubator. After incubation, the cells were washed carefully and supernatants were aspirated set to leave 50 ul/well. After washing, 100μL of infected cells were plated into wells and 100μL of Philippine crocodile serum were added. The plates were incubated for 72 hours at 37°C in the CO_2_ incubator. The contents of each well were transferred to microfuge tubes, cells were pelleted at 13,000 RPM for 5 minutes and the supernatants used for determining the HIV-1 RT levels.

#### Pre-infection interaction (cell-free HIV)

The procedure was based on the study of Dahake, R (2013). Briefly, the HIV-1 virions from co-cultured were first interacted with the Philippine crocodile serum for two (2) hour and then allowed to infect PBMCs. A volume of 10μL of HIV-1 virions were added to 10μL of Philippine crocodile serum (Human serum as a negative control) incubated for 2 hours at 37°C in a CO_2_ incubator. After 2 hours of interaction the experimental moiety-virus suspension was added to 100μL of PHA stimulated PBMCs and incubated for 2 hours in the CO_2_. After incubation, the cells were washed carefully and supernatants were aspirated set to leave 50 ul/well After washing, 100μL of infected cells were plated into wells and 100μL of Philippine crocodile serum were added. The plates were incubated for 72 hours at 37°C in the CO_2_ incubator. The contents of each well were transferred to microfuge tubes, cells were pelleted at 13,000 RPM for 5 minutes and the supernatants used for determining the HIV-1 RT levels.

#### Data Management

The results of pre-infection and post-infection assays were expressed as means ±SD in triplicates. The Percent (%) reduction of HIV-1 activity was computed using the following mathematic formula:

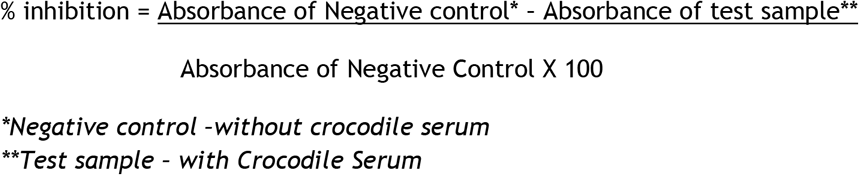

## Results and Discussion

### HIV-1 Co-culture and Cell viability

The HIV-1 RT activity after 21 days was 0.5928 ng/well. On the other hand, a baseline Philippine crocodile serum concentration of 0.5% vol/vol was used based on the study conducted by Hinay and Sarol (2018) and the cell viability results showed no cell reduction of mononuclear cells after 72 hours incubation.

### Post-infection interaction (cell-free HIV)

Table 1 shows the inhibitory activity of the Philippine crocodile serum. HIV-1 replication inhibition at 0.5% and 0.25% vol/vol concentrations were 65.68±2.93% and 69.92±0.45% respectively. This is also supported with the HIV-1 Reverse transcriptase activity of the peripheral blood mononuclear cells (Figure 1). The PBMCs interacted with crocodile serum concentration of 0.5% and 0.25% vol/vol shows a decreased HIV-1 reverse transcriptase activity of 0.1216 and 0.0954 ng/well respectively.

**Figure 1.**
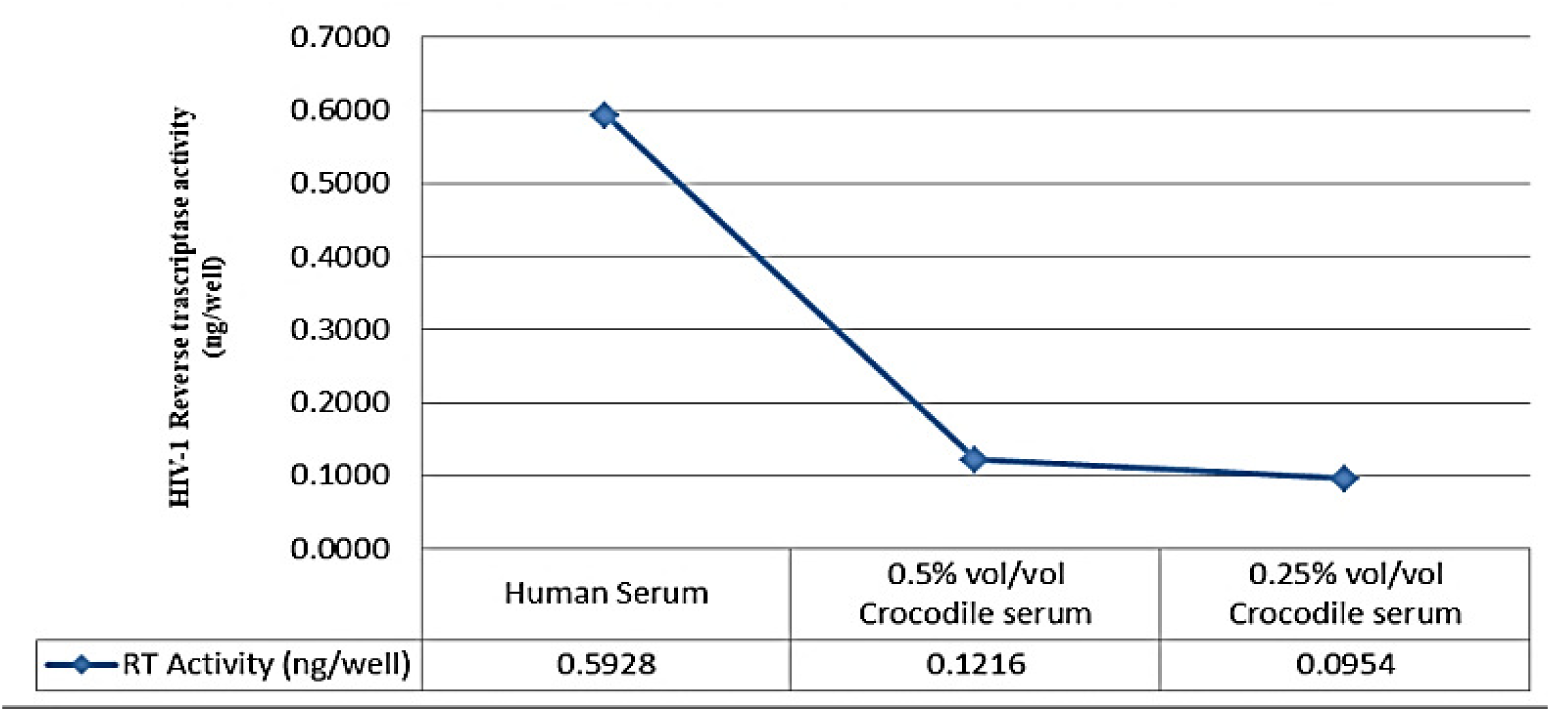
Reverse Trascriptase Activity in Post-infection interaction (cell-associated HIV)

**Table 1.** Inhibitory Activity of the Philippine Crocodile Serum in Post-infection interaction (cell-associated HIV)

| **Concentration % vol/vol** | **Post-infection**<br>**Mean±SD** | **Post-infection**<br>**Mean±SD** |
| --- | --- | --- |
| 0.50 | 65.68±2.93 | 68.61±1.67 |
| 0.25 | 69.32±0.45 | 69.95±2.24 |

Unexpectedly, an inverse relationship was observed between crocodile serum concentration and the anti-HIV-1 Reverse Transcriptase activity. This means that the inhibitory activity decreases as the Philippine crocodile serum concentration increases. This is similar to the previous study of Hinay and Sarol (2018), were the result could be explained by the presence of an “inhibitor of the inhibitor” in the crocodile serum. The best candidate inhibitor present in the crocodile serum with putative inhibition activity to HIV-1 is the Antimicrobial Peptides (AMPs) (Finger & Isberg, 2012). This is also supported by the study of Hinay and Sarol (2018) where Philippine crocodile serum concentration showed *in vitro* inhibition of HIV-1 Reverse transcriptase by high as 92.93±0.72% at 0.5 vol/vol % (Hinay and Sarol, 2018). Moreover, another putative component of the Philippine crocodile serum as described by Okada and colleagues provided a good support that the complement fragment C5a, an anaphylotoxin and a small peptidase product during complement activation, has a 30% HIV-1 Reverse transcriptase inhibitory activity (Okada, et al., 2011). The inhibition of the HIV-1 virion interacted with the Philippine crocodile serum indicates that the inhibition occurs either direct interaction to viral enzymes reverse transcriptase and or protease which are key viral enzymes of HIV-1 replication.

### Pre-infection interaction (cell-free HIV)

The potential of the Philippine crocodile serum as inhibitor against HIV-1 replication was determined and the findings demonstrated a significant inhibition of HIV-1 replication prior to interaction to PBMCs. Table 1 shows the inhibitory activity of the Philippine crocodile serum in pre-infection interaction. Reverse Transcriptase inhibition at 0.5% and 0.25% vol/vol concentrations inhibited 68.61±1.67% and 69.95±2.24% respectively. This is also supported with the HIV-1 Reverse transcriptase activity of the peripheral blood mononuclear cells (Figure 2). The PBMCs interacted with crocodile serum concentration of 0.5% and 0.25% vol/vol with an HIV-1 reverse transcriptase activity of 0.1005 and 0.10054 ng/well respectively. With this the Philippine crocodile serum have both direct virucidal effects to HIV-1 virion or the crocodile serum inhibits the viral entry subsequently results through inhibition of fusion of the viral envelope with the host cell membrane (Ray and Doms, 2006). Possible crocodile serum component such as Alpha-1 antitrypsin (AAT) which is an acute phase reactant and the most abundant circulating serine protease inhibitor can be a putative HIV-1 inhibitor (Shapiro, Pott, & Ralston, 2001). Shapiro and colleagues (2001) describe Alpha-1 antitrypsin (AAT) in their study as a natural HIV-1 antagonist. In their study the inhibitory activity of the Alpha-1 antitrypsin was observed in the induction of latent virus from chronically infected cells, viral infection and replication in PBMC, attachment CCR-5-cell assay and infection and replication in whole blood. Merchant ME and colleagues (2005) described the effectiveness of alligator serum against the laboratory strain of Human immunodeficiency virus type-1 using a cell culture-based assay. The results exhibited potent anti-HIV activity at 20-and 64-fold dilutions with 100 and 89% reduction in cytopathic effects, respectively. The study also showed that the higher concentration of the serum was difficult to assess due to inherent toxicity to the cell culture used in the study (Merchant, et al., 2005). In addition, the results suggest that both the antiHIV-1 activity and cell toxicity were mediated by serum complement activity as heat treatment of the serum destroyed both effects. Like human complement activity, alligator serum complement is sensitive to heat inactivation (Merchant, Thibodeaux, Loubser, & Elsey, 2004).

**Figure 2.**
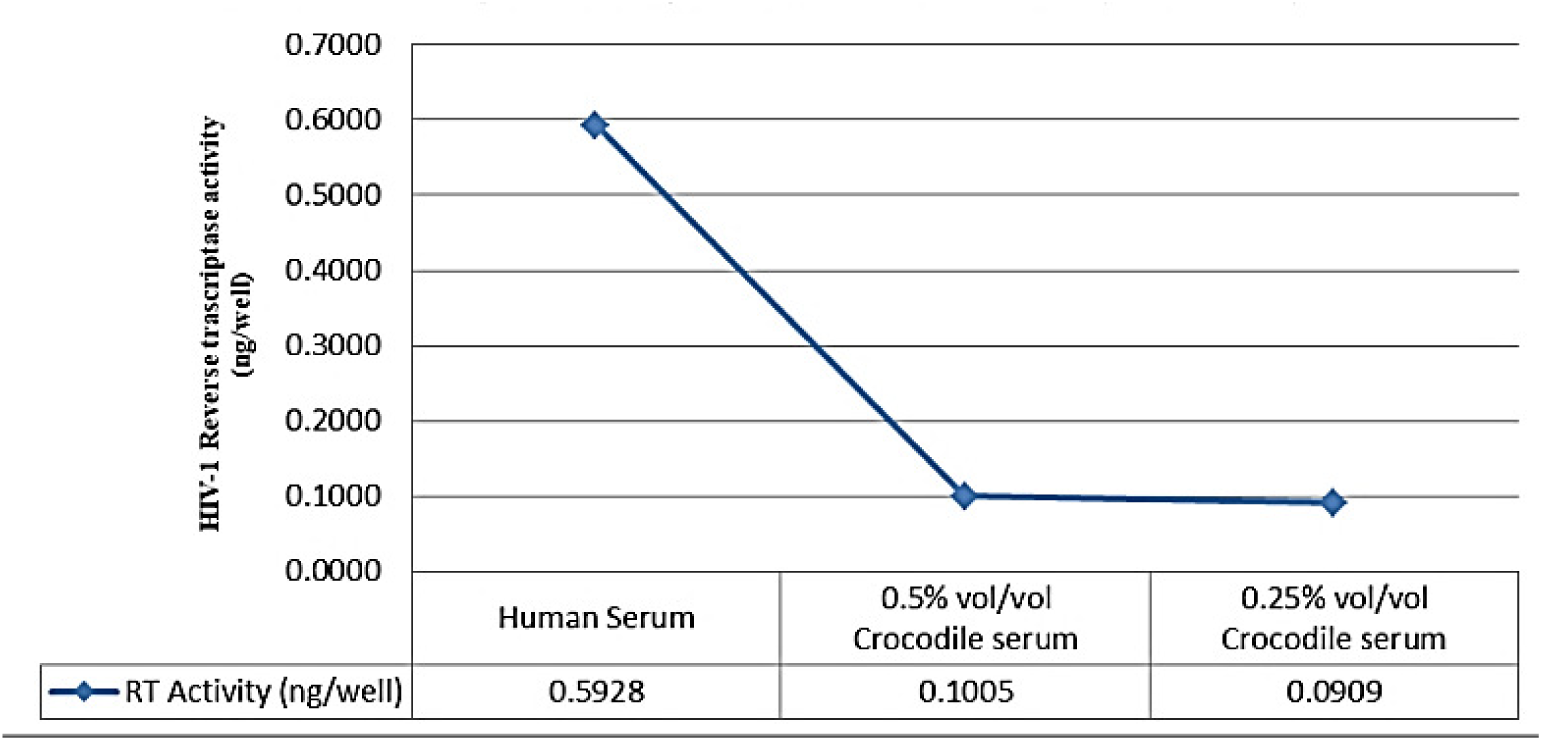
Reverse Trascriptase Activity in Pre-infection interaction (cell-free HIV)

These findings are quite significant since the Philippine crocodile serum concentration of 0.5% and 0.25% vol/vol seems to work in both pre- and post-infection interactions and hence suggest that the effect may be mediated by a direct action on the virus particle. Perhaps it blocks binding or uptake to cells or causes immediate particle lysis.

## CONCLUSION

The results showed that the HIV-1 Reverse Transcriptase activity was inhibited by 65.68±2.93% and 69.92±0.45% in Philippine crocodile serum concentration of 0.5% and 0.25% vol/vol respectively in post-infection interaction and 68.61±1.67% and 69.95±2.24% in Philippine crocodile serum concentration of 0.5% and 0.25% vol/vol respectively in pre-infection interaction. As has been noted, the 0.5% and 0.25% vol/vol concentration of the Philippine crocodile serum effectively regulates the HIV-1 replication in both pre- and post-infection interactions. With this, the study recommends to identify and characterize the active compounds in the Philippine crocodile serum and develop a novel source of HIV-1 replication inhibitors.

## ACKNOWLEDGEMENT

This work was supported and funded by the Department of Science and Technology–Regional Health Research and Development Consortium XI (*RHRDC XI*) and Davao Crocodile Park, Davao City.

## AUTHORS CONTRIBUTION

Alfredo A. Hinay Jr., Nelyn Mae T. Cadotdot and Marilou V. Tablizo designed and carried out the experiment and Alfredo A. Hinay Jr prepared the manuscript.

